# Sea anemone MACPF proteins demonstrate an evolutionary transitional state between venomous and developmental functions

**DOI:** 10.1101/2023.10.30.564692

**Authors:** Joachim M. Surm, Morani Landau, Yaara Y. Columbus-Shenkar, Yehu Moran

## Abstract

Gene duplication is a major force driving evolutionary innovation. A classic example is generating new animal toxins via duplication of physiological protein-encoding genes and recruitment into venom. While this process drives the innovation of many animal venoms, reverse-recruitment of toxins into non-venomous cells remains unresolved. Using comparative genomics, we find members of the Membrane Attack Complex and Perforin Family (MACPF) have been recruited into venom-injecting cells (cnidocytes), in soft and stony corals and sea anemones, suggesting that the ancestral MACPF was a cnidocyte expressed toxin. Further investigation into the model sea anemone *Nematostella vectensis,* reveals that three members have undergone *Nematostella*-specific duplications leading to their reverse-recruitment into mesoendodermal cells. Furthermore, simultaneous knock-down of all three mesoendodermally-expressed MACPFs leads to mis-development, supporting that these paralogs have non-venomous function. By resolving the evolutionary history and function of MACPFs in *Nematostella*, we provide the first proof for reverse-recruitment from venom to organismal development.

**Significance statement:** In this study, we reveal how a gene can gain a new function, even from a most unexpected origin. Specifically, we report that in the last common ancestor of corals and sea anemones a member of the Membrane Attack Complex and Perforin Family (MACPF), which is commonly associated with the immune system, was recruited into venom-injecting cells called cnidocytes. Using the sea anemone *Nematostella vectensis* we find repeated gene duplication has occurred leading to the new copies adopting divergent functions including being retained in cnidocytes but also recruited into non-venomous mesoendodermal cells. Furthermore, when we deplete *Nematostella* of mesoendodermally-expressed MACPFs we disrupt normal embryonic development, supporting that these copies have indeed been recruited from venom into the developmental plan.

## Introduction

In recent years, the starlet sea anemone *Nematostella vectensis* has been developed into a model system in the study of evolutionary developmental biology (1). This is due to *Nematostella* being a member of the ancient phylum Cnidaria (sea anemones, jellyfish, corals and hydroids), that diverged from Bilateria more than 600 million years ago, yet sharing a striking level of conservation in genomic features to bilaterians, such as synteny and gene content (2–6). In addition, *Nematostella* has established itself as a model to unravel the evolution and function of genes due to its ability to complete a full life cycle in the lab, its fully sequenced chromosomal-level genome and the development of new tools for its genetic manipulation, such as transgenesis methods, gene knockdowns and knockouts (7, 8). While this has been essential in unraveling characteristics of the cnidarian-bilaterian common ancestor, it has also revealed key insights into the evolution of novel innovations, in particular the evolution of venom (9–12).

Members of Cnidaria are venomous and employ specialized organelles called nematocysts as miniature venom injectors (13). These highly complex intracellular biological structures are composed of various protein polymers that are packed in stinging cells called nematocytes, then, upon specific signals, the nematocysts discharge at remarkable speed and puncture their target (13, 14). Among the list of cnidarian model systems, which is continuously expanding, *Nematostella*’s venom system is arguably the most well-studied (12). These works have revealed key insights into the molecular mechanisms underlying the nematocyte developmental origin (15, 16), such as nematocytes originating from a common neurosecretory progenitor, to their diversity, such as connecting single transcription factor (NvSox2) which can switch between two alternative stinging cell fates (17). Genetic manipulation techniques have also been crucial in characterizing toxin genes and their novel cell types (18–20) as well as resolving long-standing evolutionary theories related to venom biology such as resolving their impact on fitness (21).

In addition, these genetic tools have also been valuable in tracking the genesis of toxins from genes with non-venom functions, for example the recruitment of conserved sea anemone neuropeptide into nematocyte following a *Nematostella*-specific gene duplication event (22). This process is known as ‘recruitment’ and it is assumed that many toxins originate from gene duplication of proteins that carry physiological roles, with their new paralogs undergoing neofunctionalization to become a toxin (23). Originally, recruitment into venom was described as a one-way process, where physiological genes transform into toxins, however, a previous study in squamate reptiles hinted that this may indeed be a two-way process where “reverse recruitment”, i.e., the transformation of a venom protein back into a physiological non-venom protein, can also occur (24). However, reverse recruitment has yet to be proven experimentally.

In this work, we studied members of the membrane attack complex and perforin family (MACPF) in *Nematostella* (we refer to these as MAC). This family includes β-pore former toxins (PFTs) that are found in a wide variety of organisms from bacteria to mammals and are mostly employed for lysing cells by generating pores in their membranes (25). MACPFs were found to be toxins in the nematocysts of two sea anemones, *Actineria villosa* and *Phyllodiscus semoni* (26–29). Using comparative genomics and phylogenetics, as well as interrogating the publicly available cell atlases, we find that members of MACPF were recruited into the nematocytes of the last common ancestor of Anthozoa (soft corals, stony corals and sea anemones). We find that following two rounds of lineage-specific duplications in *Nematostella*, three additional MACPF paralogs were recruited into mesoendodermal cells strongly suggesting that they carry non-venomous functions. This is further supported by evidence that depleting these three mesoendodermal MACPFs interferes with normal development in *Nematostella*. Additionally, two of these mesoendodermal MACPF paralogs still retain some weak expression in stinging cells and represent a “transitional form” between a toxin and a non-venom protein-encoding gene. These findings are the first experimental proof of the reverse-recruitment of venom and highlight the power of gene duplication in the rapid evolution of molecular innovation.

## Results

### Evolutionary history of the MACPF family across Anthozoa

Using a phylogenetic framework, we investigated the distribution of MAC genes across Anthozoa (**Fig. 1, Table S1**). The maximum-likelihood tree reveals two distinct clades, one clade including only sea anemones and the other including soft corals, stony coral and sea anemones. Specifically, we find in the *N. vectensis* genome seven sequences that encode for proteins composed of a signal peptide and a single MACPF domain, which we named NveMAC1 through 7 (**Fig. 1**). Similarly, we find that the MAC gene family has undergone an amplification in other Hexacorallia genomes investigated to varying degrees with four copies in *Stylophora pistillata*, four copies in *Acropora millepora* two copies in *Actinia tenebrosa*, 14 copies in *Exaiptasia diaphana* and six copies in *Scolanthus callimorphus*. Within both clades, the broad clustering is consistent with anthozoan phylogeny, with Octocorallia (soft corals) being the most diverged compared to Hexacorallia (which includes stony corals and sea anemones) and within the sea anemones clade, the superfamilies Metridioidea and Actinioidea cluster together and species from Edwardsioidea are the most diverged. Beyond this broad clustering, we see repeated evidence of species-specific clustering, in Hexacorallia, suggesting lineage-specific duplications are underlying much of the evolution of this gene family. For example, we find that *Nematostella* (Nve) MACs 1-4 cluster together, in *E. diaphana* MACs 5-14 cluster together and all *S. pistillata* and *A. millepora* sequences cluster in a species-specific manner.

**Fig. 1.**
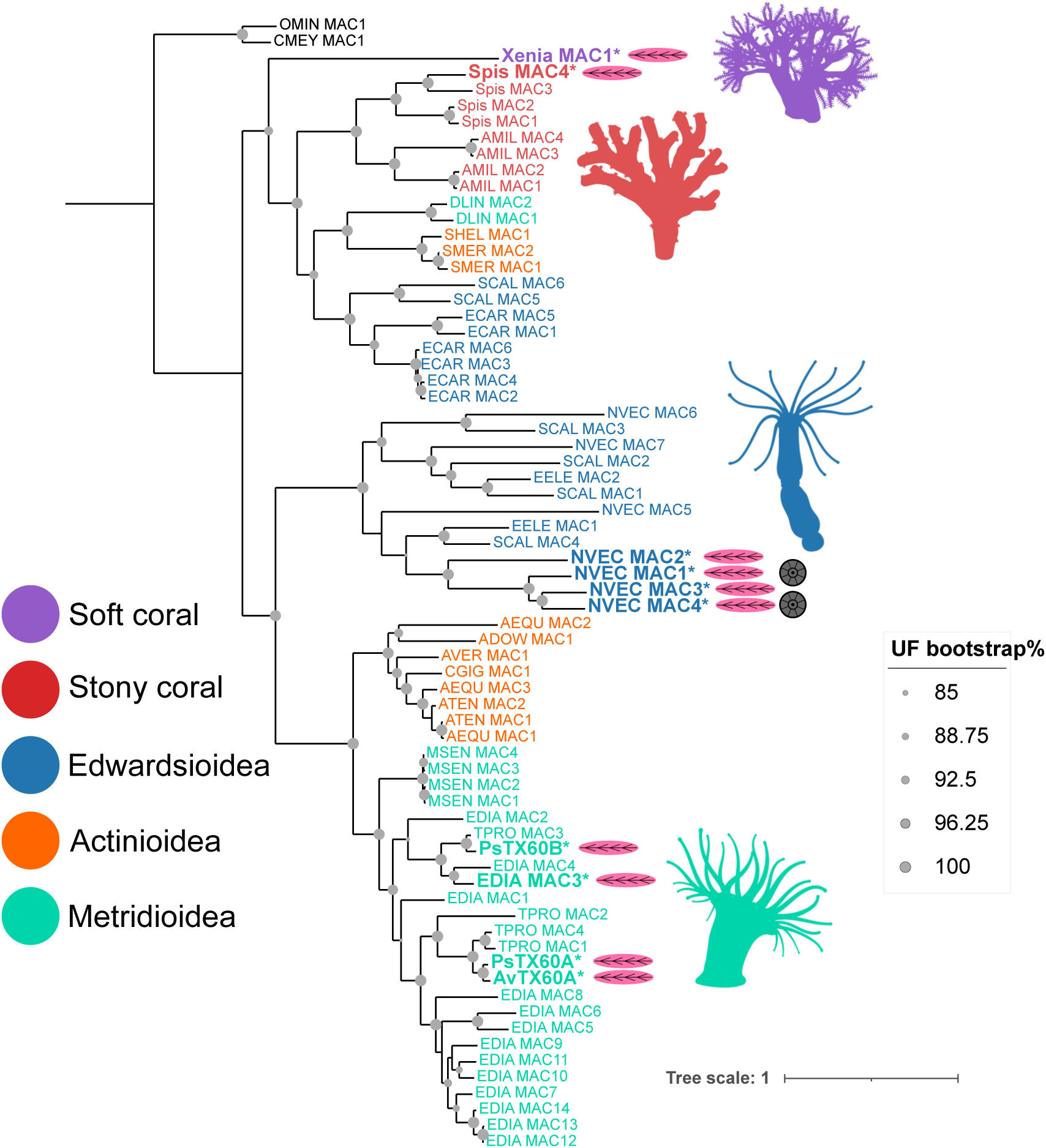
Phylogeny of MACPF genes across Anthozoa. (A) Maximum-likelihood tree of the MACPFs in anthozoans. The sequences found in nematocytes appear in bold with a magenta nematocyte cartoon. Sequences found in the Endo-atlas of *Nematostella* (30) also contain a grey cartoon representing the mesoendodermal segments. Ultrafast bootstrap values and SH-like approximate likelihood ratio test above 85 are shown as a grey circles.

While we report lineage-specific duplications among *Nematostella* MACs 1-4, we also find that NveMACs 5 and 6 cluster with *S. callimorphus* (Sca) MACs 4 and 3, respectively, and NveMAC7 clusters with ScaMAC1 (**Fig. 1**). These sequences that cluster together between *N. vectensis* and *S. callimorphus* also share chromosomal macrosynteny (**Fig. 2, Table S2**). This strongly supports that NveMACs 5-7 are likely ancestral sequences, with MACs 1-4 evolving via *Nematostella*-specific duplications. Furthermore, sea anemone MACs from the three superfamilies investigated can be found in both MAC clades. The clade containing only sea anemone sequences, however, includes sequences known to be a part of the sea anemone venom profiles. Specifically, the sequences from *P. semoni* (PsTX60A and PsTX60B) and *A. villosa* (AvTX60B) that have been isolated from nematocytes and shown to be toxic to shrimp and hemolytic towards sheep red blood cells (26–29). Using Alphafold, analysis of the predicted structure of NveMAC 1-4 with the predicted structure ofPsTX60A and PsTX60B showed broad overlap among all sequences, suggesting that potentially NveMAC 1-4 might also have hemolytic activity (**Fig. S1**). This hinted to us that these sequences may therefore also play a role in the venom composition of other sea anemones including *Nematostella*.

**Fig. 2.**
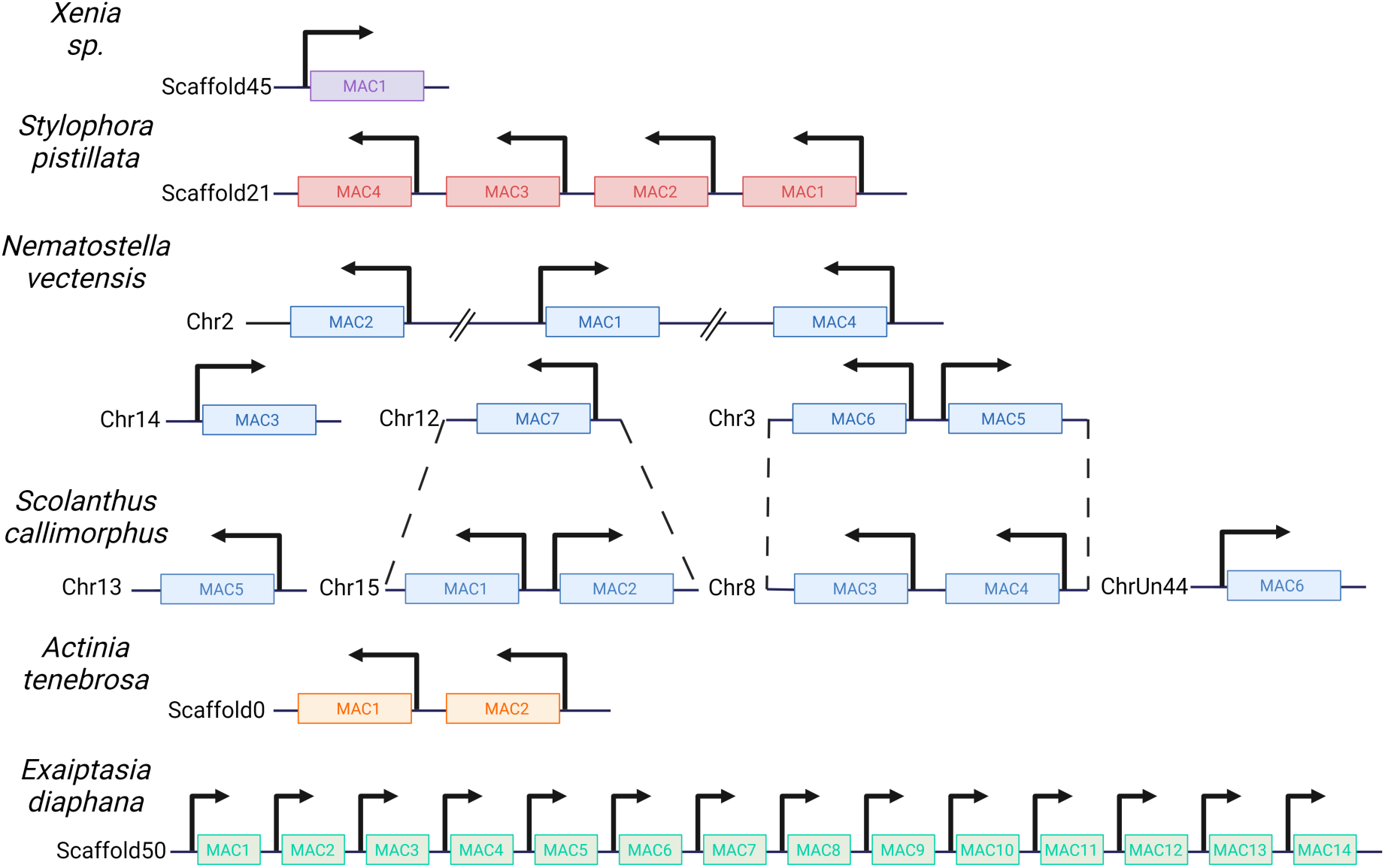
Schematic representation depicting the MACPF gene cluster and gene synteny among Anthozoans. Dashed lines between *Nematostella vectensis* and *Scolanthus callimorphus* represents a cluster that shares macrosynteny.

### Spatiotemporal expression of MACPF in *Nematostella*

The recent advancements in single-cell sequencing in non-model organisms have allowed the establishment of cell atlases of multiple different anthozoans including *Xenia sp.*, *S. pistillata*, *E. diaphana* and *Nematostella* (15, 31–33). Strikingly, we find at least one MAC is expressed in nematocytes from each of the anthozoan cell atlases (**Fig. 1, Table S1)**. In *Nematostella*, which arguably has the most comprehensive cell-atlas coming from detailed scRNA-seq across development, we find that NveMACs 1-4 have expression in nematocytes. To complement this, we investigated bulk RNA-seq of the *Nematostella* transgenic line expressing NvNcol3::memOrange2, a nematocyte marker (20, 34). We find that all *Nematostella* MACs, except NveMAC5, are upregulated in nematocytes, further supporting that *Nematostella* MACs are expressed in nematocytes, including the more ancestral sequence NveMAC6 and 7. Interestingly, in *Nematostella* scRNA-seq, NveMAC5 can be seen to be expressed in neurons. This is further supported by bulk RNA-seq data coming from the *Nematostella* transgenic line expressing NvElav1::memOrange, a neuronal marker (35, 36), with NveMAC5 significantly upregulated in neurons. While NveMAC5 and NveMAC6 are genomic neighbors and positioned head-to-head, a feature that is often associated with genes sharing the same protomers and cis-regulatory genetic information, *Nematostella* Chip-seq data (**Fig. S2**), suggests that these two genes share different promotor regions which would allow for a divergence in gene regulation cell-type specific expression.

Finally, a recent endomesodermal-enriched scRNA-seq dataset from planula was generated to construct a 3D spatial gene expression atlas of *Nematostella*. From this dataset, we find that two MACs, MACs 1 and 4, are expressed at relatively high levels compared to the other MACs as well as all other toxins and nematocyte makers (**Table S1 and S3**). This finding suggests that these two MACs are expressed in the mesoendoderm in addition to nematocytes. This is further supported by RNA-seq following the knockout of HOX genes (ANthox1a, ANthox6a, ANthox8), as the Knockout of these important regulators of endomesodermal segmentation in *Nematostella* resulted in significant down regulation of NveMAC1 and NveMAC4 (**Table S8**). A similar pattern is observed for other genes known to be expressed in the mesoendoderm (30).

Due to these conflicting results coming from different *Nematostella* scRNA-seq datasets, we aimed to characterize the expression of NveMACs experimentally using in situ hybridization (ISH). Interestingly, of the four genes that are expressed at levels measurable by ISH, only NveMAC2 is being expressed exclusively in nematocytes, whereas NveMAC1 is expressed in mesoendodermal cells (**Fig. 3A**). To verify the specificity of the obtained ISH patterns we microinjected to *Nematostella* zygotes shRNA against NveMAC1 and showed that the staining is greatly reduced in comparison to larvae of the same age that developed from zygotes injected with control shRNA (**Fig. S3):** NveMAC1 knockdown: 3/189 stained embryos; control shRNA 145/146 stained embryos. In parallel, we explored the temporal expression of NveMACs revealing that NveMAC1 expression is restricted to a very short period, peaking at the planula stage (**Fig. 3B** and **S3, Table S4, S5**). In contrast, NveMAC2 expression is relatively stable throughout the life of *Nematostella*. Our PCA analysis of expression profiles across early development (0-240hpf) further supports this, with NveMAC1 and NveMAC2 grouping separately (**Fig. 3C**). Taken altogether, NveMAC1 and NveMAC2 have distinct spatiotemporal expression patterns, with NveMAC2 likely playing a role in *Nematostella* venom, and the relatively short time window of the expression of NveMAC1 raises the unexpected possibility that it might be involved in development.

**Fig. 3.**
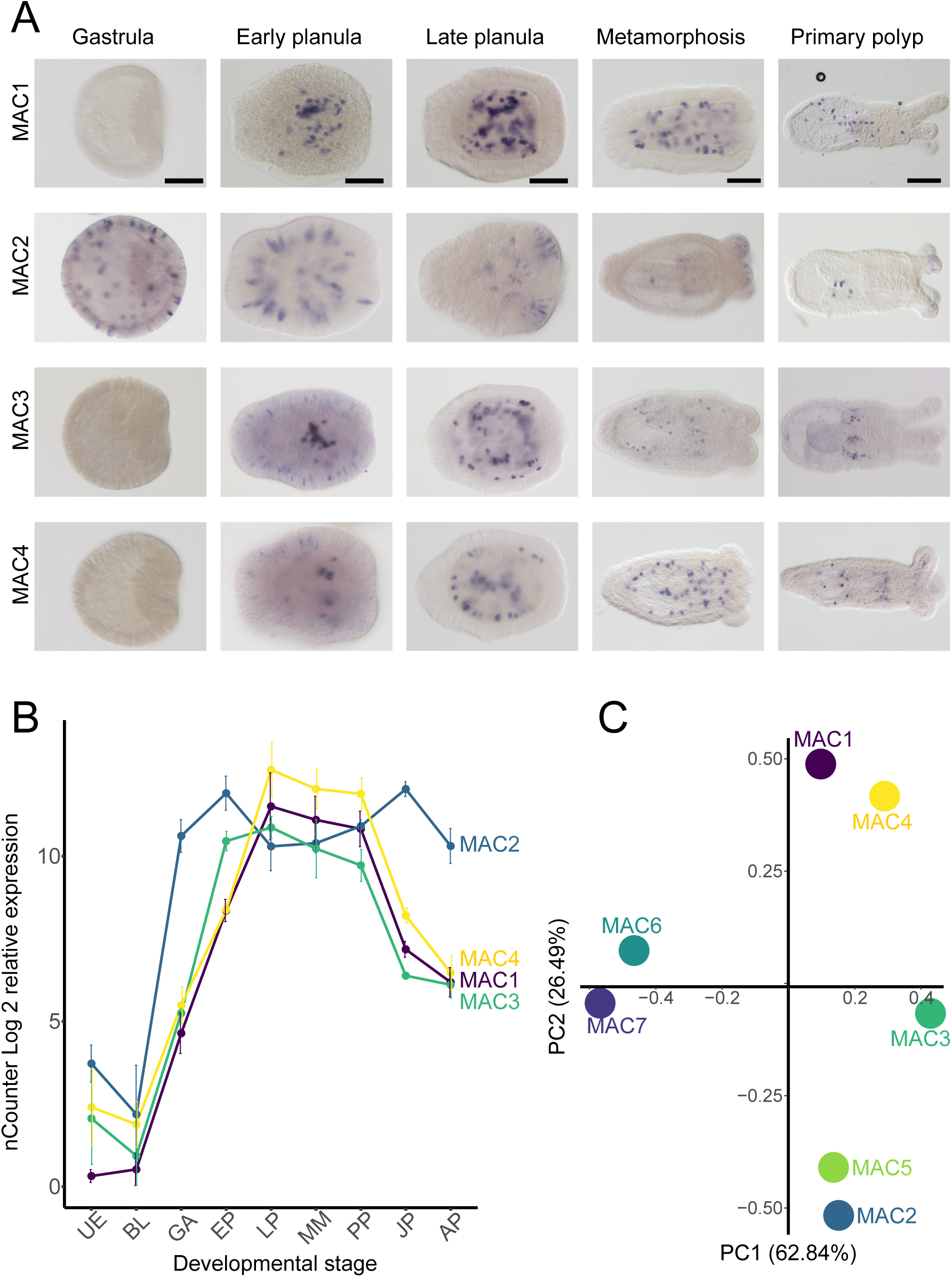
Spatiotemporal expression of MACPF-encoding genes in *Nematostella*. A) in situ hybridization of NveMACs (1–4) across embryonic development. B) Graph of RNA levels of the same MACPF genes used in ISH. Scale bar represents 100 μm. C) Principal component analysis (PCA) using temporal expression of NveMACs in *Nematostella* throughout embryonic development taken from the NvERTx database. UE = unfertilized egg; BL = blastula; GA = gastrula; EP = early planula; LP = late planula; MM = metamorphosis; PP = primary polyp; JP = juvenile polyp; AP = adult polyp

In addition, NveMAC3 and 4 present an “intermediate” expression pattern. In the early planula, they are weakly expressed in nematocytes, while strongly expressed in endodermal cells resembling those that express NveMAC1 (**Fig. 3A**). Additionally, NveMAC4 temporal expression pattern mirrors the expression pattern of NveMAC1 and is supported by our principal component analysis (PCA) which clusters NveMAC1 and 4 together (**Fig. 3B and C)**. Interestingly, NveMAC3 temporal expression clusters uniquely in between NveMAC1 and NveMAC2, further suggesting that NveMAC3 expression pattern is a combination of NveMAC1 and NveMAC2. Given the strong expression profile of NveMACs 1 3 and 4 across development and their expression in mesoendodermal cells we suspected that they have non-venomous function and may play a role in development.

### MACPF depletion interferes with *Nematostella* development

To investigate the functional role of MACs 1-4, we depleted their expression in embryos by injecting short-hairpin RNA (shRNA) targeting NveMACs specifically. Knockdown efficiency was confirmed in four-day-old planula using qPCR, revealing that all shRNA used resulted in significant knockdown efficiency (**Table S6**,>50 % knockdown and P-value <0.05) of the target NveMAC compared to animals injected with control shRNA. After confirming the knockdown efficiency, additional animals were tracked until 10 days post-fertilization (dpf). The control shRNA-injected embryos developed normally, undergoing metamorphosis and progressing into primary polyps. shRNA-injected embryos targeting specific NveMACs (1–4) also resulted in normal developments, with animals undergoing normal metamorphosis (**Table S7**).

Given the overlap of RNA expression for NveMACs 1, 3 and 4 in the mesoendoderm, we suspected that some compensation might be occurring. To test this, we co-injected shRNAs to target NveMACs 1, 3 and 4 simultaneously. We confirmed that this approach still results in a significant knockdown (**Fig. 4 A-C**, >50%; P-value <0.05) of all three NveMACs (1, 3 and 4) compared to planula injected with an equal concentration of control shRNA. Strikingly, embryos injected with a combination of shRNAs targeting MACs 1, 3 and 4 have developmental defects, with only ∼65% of 10 dpf of these animals developing into primary polyps compared to control (**Fig 4 D-F**, P-value=0.02), in which 85% of animals underwent metamorphosis into primary polyps by the same time point. Developmental defects in NveMACs 1, 3 and 4-depleted animals suggest these members of the MAC family are essential for proper *Nematostella* development.

**Fig. 4.**
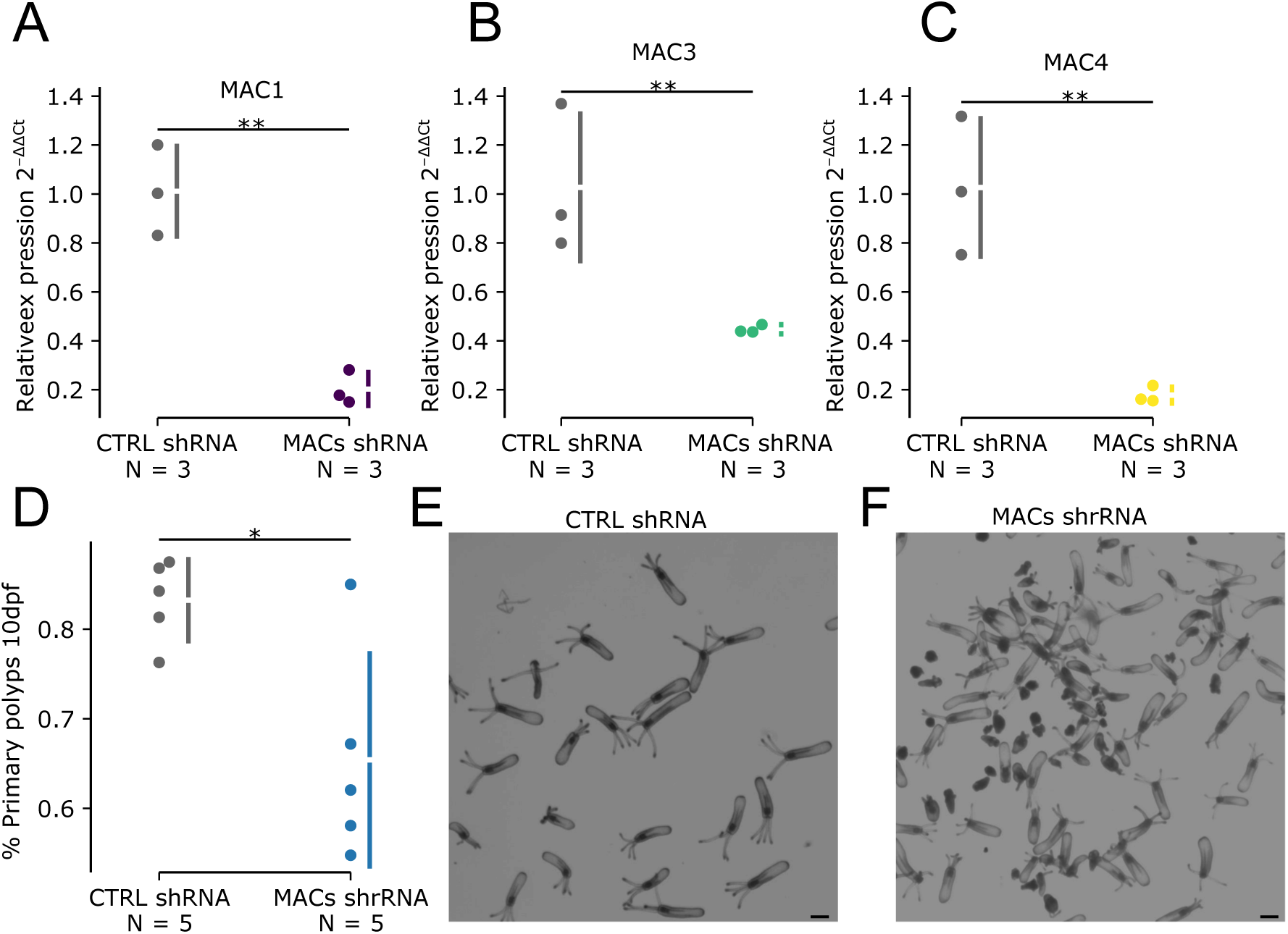
Knockdown of MACPFs in *Nematostella*. qRT-PCR for NveMACs at late planula stage after shRNA injection. The graph shows the relative foldchange in expression between control shRNA and the combined knockdown of (A) NveMAC1, (B) NveMAC3 and (C) NveMAC4, using sequence-specific shRNA. The values for the individual replicates are shown as circles. The mean difference is depicted as a dot; the 95% confidence interval is indicated by the ends of the vertical error bar. D) Quantification of normal polyps development following the injection of control shRNA or knockdown of NveMACs 1, 3, and 4 simultaneously in 10-dpf polyps. E) control shRNA, F) NveMACs 1, 3 anand 4shRNA. * P-value < 0.05, NS not significant **P-value < 0.01. Scale bar represents 200 μm

## Discussion

In this work, we unraveled the evolution of a gene family of proteins containing the MACPF domain in *Nematostella* as its role transitions from venom to development. Broadly, proteins that contain the MACPF domain are part of a superfamily of β-pore former toxins that have a large distribution, covering various groups of the tree of life, from bacteria to mammals (25, 37). The structure of MACPFs enables their function in lysing cells by generating pores in their membranes (25, 37). This ability to cause pores in cell membranes is useful for a variety of roles in eukaryotes ranging from immunity to development (38). Two prominent examples are Astrotactins from mammals and Torso-like from flies that carry important developmental roles (39, 40). Notably, some proteins that contain a MACPF domain, such as Astrotactin-2, can carry functions even if they are unable to lyse cells (37, 41).

The MACPF domain containing proteins have undergone considerable variation at the sequence level, yet their predicted 3D structure is relatively well conserved. Notably, the sequences from *P. semoni*, which are known to lyse blood cells, share this high structural similarity (26, 28). Given this conservation in structure, we suspect that all seven *Nematostella* MACs maintain the capacity to lyse cells, especially copies 1-4 which share the greatest similarity to *P. semoni* MACs. This ability to disrupt cell membranes can serve different cellular functions, including embryonic development (38). The recruitment of several MACs into developmental processes is a striking example for neofunctionalization.

It should be noted that while we see no evidence of nematocyte expression of NveMAC1 in our ISH results, the scRNA-seq analysis suggests that this gene still exhibits some expression in nematocytes. This residual expression is caught by ISH for NveMAC3 and NveMAC4, suggesting it is more profound for these genes. The very weak expression of NveMACs 1, 3 and 4 in cnidocytes compared to their mesoendodermal expression domain suggests that this is “vestigial expression” that might becoming non-functional. Furthermore, the expression of NveMAC3 and NveMAC4 both in nematocytes and in mesoendodermal cells suggests that they may be a molecular transitional state between the ancestral expression domain in nematocytes to the derived expression domain in the mesoendoderm.

The most parsimonious explanation for our findings is that the ancestral sequence of NveMACs 1, 3 and 4 likely was expressed in both nematocytes and mesoendodermal cells. We, therefore, suggest that the ancestral MAC1/3/4 gene was reverse-recruited out of venom and gaining a new function, being expressed in the mesoendodermal cells. Following duplication leading to the birth of MAC1 and MAC4, we suspect that the residual expression of NveMAC1 in nematocytes is nearly lost. This supports that MAC1 is undergoing specialization in the mesoendodermal cells. Taken together with evidence that MAC1 and MAC4 share very similar temporal expression patterns, it suggests that the specialization we see in NveMAC1 occurred after the duplication and generating MACs 1 and 4. We also suspect that specialization of MAC3 and MAC4 is also occurring with the nematocyte expression being lost, however, this loss is occurring via a more gradual process.

Finally, NveMAC5 is another member of this gene family also likely being reverse recruited but into neurons. Again, given that MAC6 and 7 are expressed in nematocytes and are more basally-branching, this suggests that MAC4 has also undergone reverse-recruitment. Interestingly, NveMAC5 andNveMAC6 are clustered and they both share macrosynteny with ScaMAC3 and ScaMAC4 which cluster together (**Fig. 1**). However, the spatiotemporal expression of the ScaMACs remains unresolved, limiting our ability to reconstruct if this reverse recruitment is shared between *Scolanthus* and *Nematostella*. While we see no residual expression of NveMAC5 in nematocytes, the temporal expression is like NveMAC2, clustered together in our PCA analysis (**Fig. 3C**). Consistent with our phylogenetic analysis (**Fig. 1**), it would suggest that the ancestral NveMAC2/5 had similar temporal expression patterns and following the duplication event, we suspect NveMAC5 was reverse-recruited into neurons from nematocytes. Furthermore, NveMAC5 and NveMAC6 are genomic neighbors and positioned head-to-head, however, *Nematostella* Chip-seq data suggests that they have different promoters which would allow them to distinct gene regulation to drive expression in neurons to nematocytes, respectively. Taken together with our results for NveMACs 1, 3 and 4, we have captured two independent events for the reverse recruitment of a venom component.

Plausibly, maintaining the ancestral nematocyte expression of NveMAC3 and NveMAC4 may not be deleterious and therefore is kept due to neutrality. This is supported by the evidence that these genes arise from more recent duplication events in which not enough time has occurred for them to lose this ancestral nematocyte expression via drift. This is consistent with previous systematic studies investigating gene duplicates in fungi: In this work, the authors show that genes involved in complex interactions, such as those essential for cell growth, are sensitive to increased gene expression noise associated with increased copy number and tend to not evolve via gene duplication (42). Contrastingly, genes that are responsive to stress and have dynamic gene expression levels tend to evolve via gene duplication. This is consistent with the function of venom-related genes, which are highly responsive to stress in *Nematostella* and have undergone significant copy-number variation across populations to meet their ecological requirements (21, 43, 44).

We find development is disrupted when MAC1, 3 and 4 are depleted simultaneously. A potential explanation for this finding is that these genes are redundant and the expression of even one of them is enough to allow normal development. Alternatively, it is tempting to speculate that these three MACs may need to oligomerize to a form protein complex. This is reasonable given that other proteins containing MACPF domains also oligomerize to form complexes, such as the human complement proteins C6-C9 which all contain a MACPF domain and assemble into the membrane attack complex (37).

Overall, our findings have uncovered the striking evolutionary history of this gene family containing a MACPF domain in Anthozoa. We reveal that gene duplication is driving the recruitment of different members of this gene family into different cell types ranging from nematocytes, neurons and mesoendodermal cells. We have also discovered that members that have undergone more recent duplication events are currently going through a transitional state, where they have gained expression in a new cell type while still maintaining residual expression from their ancestral copy. This transitional state is an exciting discovery as piecing together the exact evolutionary process that leads to genes deriving their function is extremely difficult. The mechanistic basis for the expression of these genes in multiple spatial domains remains to be discovered.

## Methods

### Comparative genomics and phylogenetics

We analyzed transcriptomes from 10 sea anemone species, spanning three of the five Actiniarian superfamilies (Actinioidea, Edwardsioidea, and Metrioidea). These transcriptomes that were sampled from either multiple tissues or tentacles were downloaded from NCBI SRA using FASTQ-DUMP in the SRA toolkit. Raw reads retrieved were assessed for quality and trimmed using Trimmomatic (45). Trinity was used to assemble transcriptomes de novo from the filtered raw reads (46, 47). BUSCO (v4) was used to validate the quality and completeness of the transcriptomes (48). We predicted open-reading frames from each transcriptome using ORFfinder (https://www.ncbi.nlm.nih.gov/orffinder/) and performed BLASTp (E-value 1e-05) using sea anemones MACPF toxins as queries. Sequences retained were used to determine the presence of a signal peptide using SignalP (v5.098) as well as a single MACPF Pfam domain (PF01823.22). A similar pipeline was also performed to identify MACPFs across Anthozoa using genomes from *Xenia* sp., *A. millepora*, *S. pistillata*, *N. vectensis*, *S. callimorphus, A. tenebrosa*, *Actinia equina* and *E. diaphana (*CC7). Sequences were then manually curated, and those including large insertions or deletions were removed.

The refined list of full-length MACPFs was used for phylogenetic analyses to determine the reconstruct its evolutionary history across Anthozoa. Protein sequences were aligned using MUSCLE in MEGA 11 (49). Protein alignments were imported into IQ-TREE and the best fit of protein model evolution was determined using ModelFinder (50). Using the Bayesian information criterion, a WAG+I+G4 model was selected as the best-fit model of protein evolution. Phylogenetic trees were generated from alignments using 1,000 ultrafast bootstrap iterations and the SH-aLRT test (51). The tree was visualized using Interactive Tree Of Life (52).

Homologous chromosomes were found among the genomes to determine if anthozoan MACs show any evidence of macrosynteny among anthozoans. This was achieved first by identifying 1103 single-copy orthologs using OrthoFinder (53) with the predicted proteomes annotated from anthozoan genomes that were generated using long-read sequencing technology. This included *A. millepora*, *N*. *vectensis*, *S. callimorphus* and *A. tenebrosa*. The chromosomal locations for the single-copy orthologs were then compared to generate a macrosynteny map of chromosomes among the genomes. For the remaining genomes, we extracted all predicted proteins from the same scaffolds containing MACs and performed BLASTp (1e-5) against *N. vectensis* proteins and counted their chromosomal location in the *N. vectensis* genome. For *E. diaphana*, scaffold50 was found to contain all MACS used in this study and was downloaded from Reef Genomics (http://aiptasiav2.reefgenomics.org/). Proteins were then predicted using the FGENESH (54) online server (http://www.softberry.com/berry.phtml).

### Sea anemone culture

*Nematostella* embryos, larvae and juveniles were grown in 16 ‰ sea salt water at 22 °C. Adults were grown in the same salinity but at 17 °C. Polyps were fed with *Artemia salina* nauplii three times a week. Induction of gamete spawning was performed according to a published protocol (55). The gelatinous egg sack was removed using 3 % L-cysteine (Merck Millipore, Burlington, MA) and followed by microinjection of shRNAs. All *N. vectensis* individuals used in this study belonged to the common lab strain originating from Rhode River MD (1).

### nCounter analysis

Total RNA from different developmental stages of *Nematostella* was extracted as previously described (18). Briefly, RNA was extracted using Tri-Reagent (Sigma-Aldrich, St. Louis, MO) according to manufacturer’s protocol, treated with Turbo DNAse (Thermo Fisher Scientific, Waltham, MA) and then re-extracted with Tri-Reagent. RNA quality was assessed on Bioanalyzer Nanochip (Agilent, Santa Clara, CA). Each sample was prepared from hundreds of specimens in order to normalize for any individual variation. Gene expression of MACs was analyzed using the nCounter platform (NanoString Technologies, Seattle, WA; performed by Agentek Ltd., Israel) as previously described (18), using technical triplicates, each made from a different batch of RNA sample. For each MAC transcript tested, we used two probes each. Normalization we performed using the geometric mean of the expression levels of five reference genes with stable expression across development (18).

### shRNA Generation and KD Experiments

Two shRNA precursors for each MAC gene was designed as previously described (56, 57). Reverse complement sequence of shRNA precursors were synthesized as DNA ultramer oligo by Integrated DNA Technologies (Coralville, IA), mixed with T7 promoter primer in 1:1 ratio in a final concentration of 25 µM, denatured at 98 °C for 5min, and cooled to 24 °C. shRNAs were synthesized with AmpliScribe T7-Flash Transcription Kit (Epicentre, Charlotte, NC) for 15 h followed by 15 min treatment with 1 µl of DNase I. The in vitro transcribed products were purified using the Quick-RNA Miniprep Kit (Zymo Research, Irvine, CA). shRNAs were used for microinjection at concentrations ranging from 400-1200 ng/ul. ∼100 injected planula (4 dpf) were flash frozen in liquid nitrogen, and stored at −80 °C and used for downstream qPCR analysis. MAC1-4 were first targeted individually and then MAC1, 3 and 4 were targeted simultaneously by combining three validated shRNA’s each targeting MAC1, 3 or 4 specifically.

### Reverse-Transcription Quantitative PCR

To quantify the knockdown efficiency of our shRNAs we analyzed the expression levels of MACs using by reverse-transcription quantitative PCR (RT-qPCR). A minimum of three biological was used for each shRNA or combination of shRNAs. First, RNA was extracted from injected embryos following the same protocol as previously described (57). 500 ng of RNA was converted into cDNA in a 20 μl reaction. cDNA was constructed using iScript cDNA Synthesis Kit (Bio-Rad, Hercules, CA) according to the manufacturer’s protocol. Real-time PCR was prepared with Fast SYBR Green Master Mix (Thermo Fisher Scientific) on the StepOnePlus Real-Time PCR System v2.2 (ABI, Thermo Fisher Scientific). The expression levels of tested genes were normalized to previously validated housekeeping gene (18), and the relative gene expression was calculated using the 2ΔΔCt method. The significance level was calculated by two-tailed Student’s t-test to ΔCt values for each of the pairwise comparisons to control shRNA.

### Assessment of phenotype following KD of MACs

We injected either a specific shRNA or a combination of shRNAs. Control shRNA were injected in parallel at an equal concertation and morphology was tracked for injected animals until 10 dpf. Each experiment consisted of at least three biological replicates. For the combinations of shRNAs to target MAC1, 3 and 4, 400 ng/µl was injected of each specific shRNA, whereas 1200 ng/µl of control shRNA was injected in parallel. After 10 dpf, animal physiology was visualized under an SMZ18 stereomicroscope equipped with a DS-Qi2 camera (Nikon, Tokyo, Japan).

### In situ hybridization (ISH)

In situ hybridization (ISH) was performed as previously described (58). Embryos older than 4 days were treated with 2 u/µl T1 RNAse (Thermo Fisher Scientific) after probe washing in order to reduce background. Stained embryos and larvae were visualized with an Eclipse Ni-U microscope equipped with a DS-Ri2 camera and an Elements BR software (Nikon). For each gene, at least 20 individuals from each developmental stage were tested. The specificity of the NveMAC1 probe was confirmed by performing ISH on 4 dpf animals that were injected with either control shRNA or shRNA to knockdown MAC1. This was repeated and the ratio of stained animals was compared.

### Meta analysis of bulk RNA-seq scRNA-across

We performed a comparative analysis using previously published RNA-seq data of two reporter lines and three mutant lines. Data from both reporter lines were generated from *Nematostella* primary polyps that express a fluorescent transgene, under either the promoter of NvNcol3::memOrange2, a nematocyte maker (20, 34), or NvElav1::memOrange, a neuronal marker (35, 36). The data from mutant lines consisted of HOX homozygote knockout lines (ANthox1a, ANthox6a, ANthox8) which play an important role in regulating endomesodermal segmentation in *Nematostella* (30). Raw reads were downloaded from the Sequence read archive (NvNcol3::memOrange2: PRJEB40304, NvElav1::memOrange PRJEB36771, HOX mutants: PRJNA727015) Raw reads were trimmed and quality filtered by Trimmomatic. Reads were mapped to a modified *Nematostella* genome. Mapping was performed using STAR and the gene counts quantified using RSEM. Differential expression analyses were performed using scripts from Trinity using both DESeq v2.139 and edgeR v3.1675. Gene models used in all downstream analyses were from previously published annotations (59). Differentially expressed genes were defined by FDR < 0.05 and fold change ≥ 2. Genes identified by both methods were considered as differentially expressed. Biological replicates were quality-checked for batch effect using sample correlation and principal component analysis.

### Structural predictions

AlphaFold2 was used to model the structure of *P. semoni* MACs and NveMACs 1-7 (60). Top-ranked AlphaFold2 models for each MAC was used as a query to search the AlphaFold/UniProt50 and PDB database using the Foldseek webserver in TM-align mode (61). Predicted structure figures were generated using PyMOL version 2.4.0 (Schrödinger, LLC).

## Author contributions

Conceptualization: JMS, YM

Methodology: JMS, ML, YYC-S

Visualization: JMS, YYC-S

Funding acquisition: JMS, YM

Supervision: JMS, YM

Writing – original draft: JMS, YM

Writing – review & editing: All authors

