## Supplemental Material for "Sea anemone MACPF proteins demonstrate an evolutionary transitional state between venomous and developmental functions"

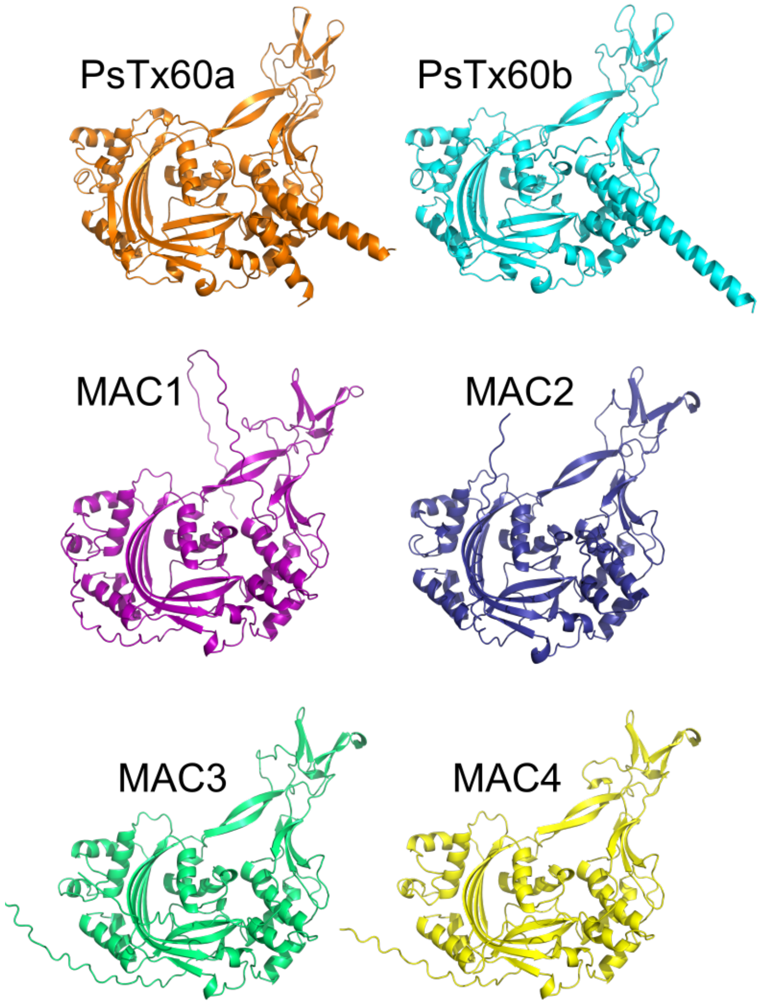


**Fig. S1.** Alphafold Predicted structures of MACPF-encoding proteins in *Phyllodiscus semoni* (PsTx60a and PsTx60b) and *Nematostella vectensis* (MAC1-4).

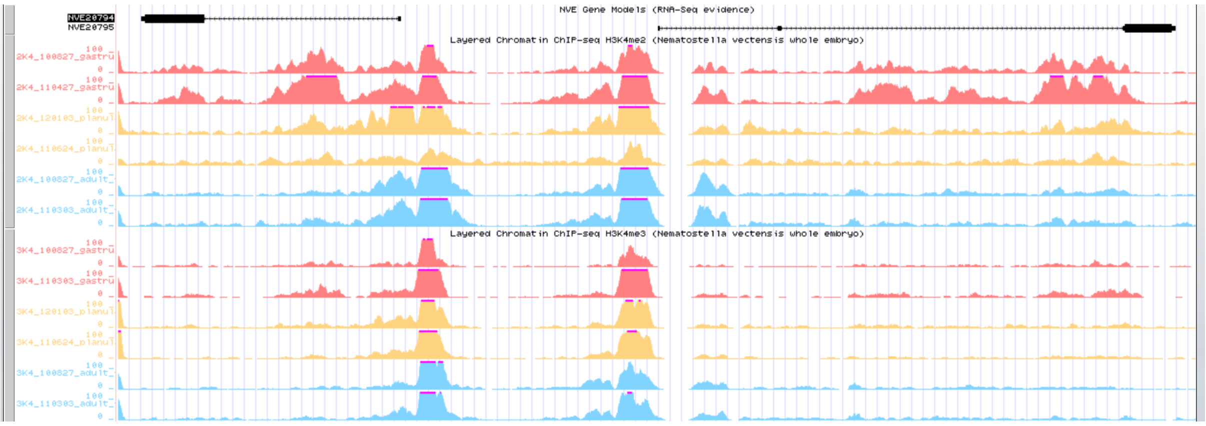


**Fig. S2** Chip-seq mapping across NveMAC5 (NVE20795) and NveMAC6 (NVE20794).

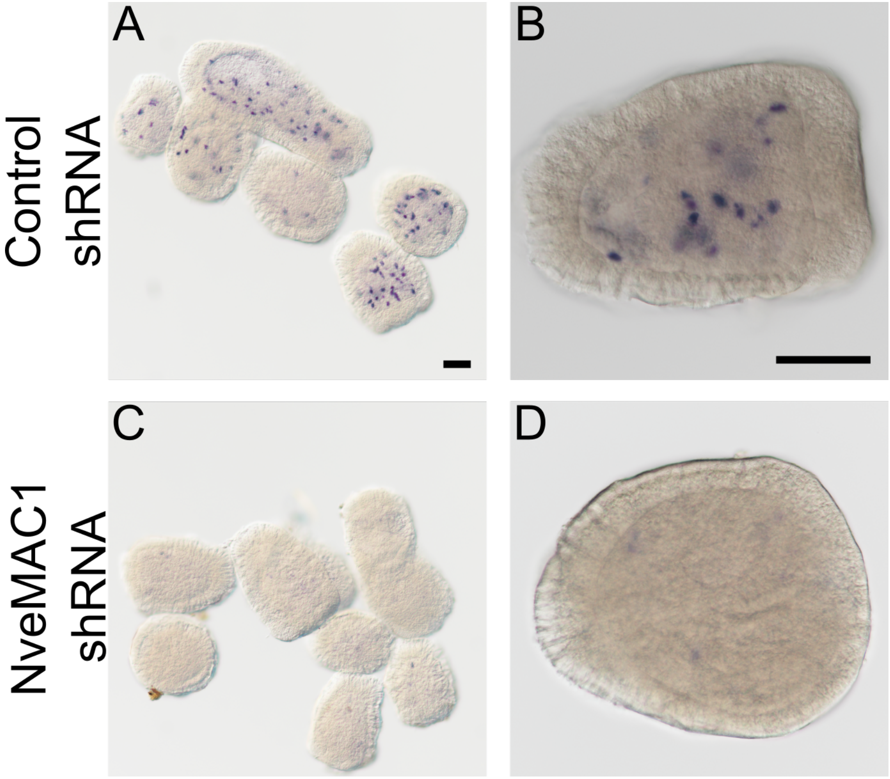


**Fig. S3.** In situ hybridization probe specificity validation. Multiple (A) and an individual (B) 4dpf *Nematostella* stained using MAC1 probe following injection of control shRNA. Multiple (C) and an individual (D) 4dpf *Nematostella* stained using MAC1 probe following injection of shRNA targeting NveMAC1. Scale bars, 100 µm.

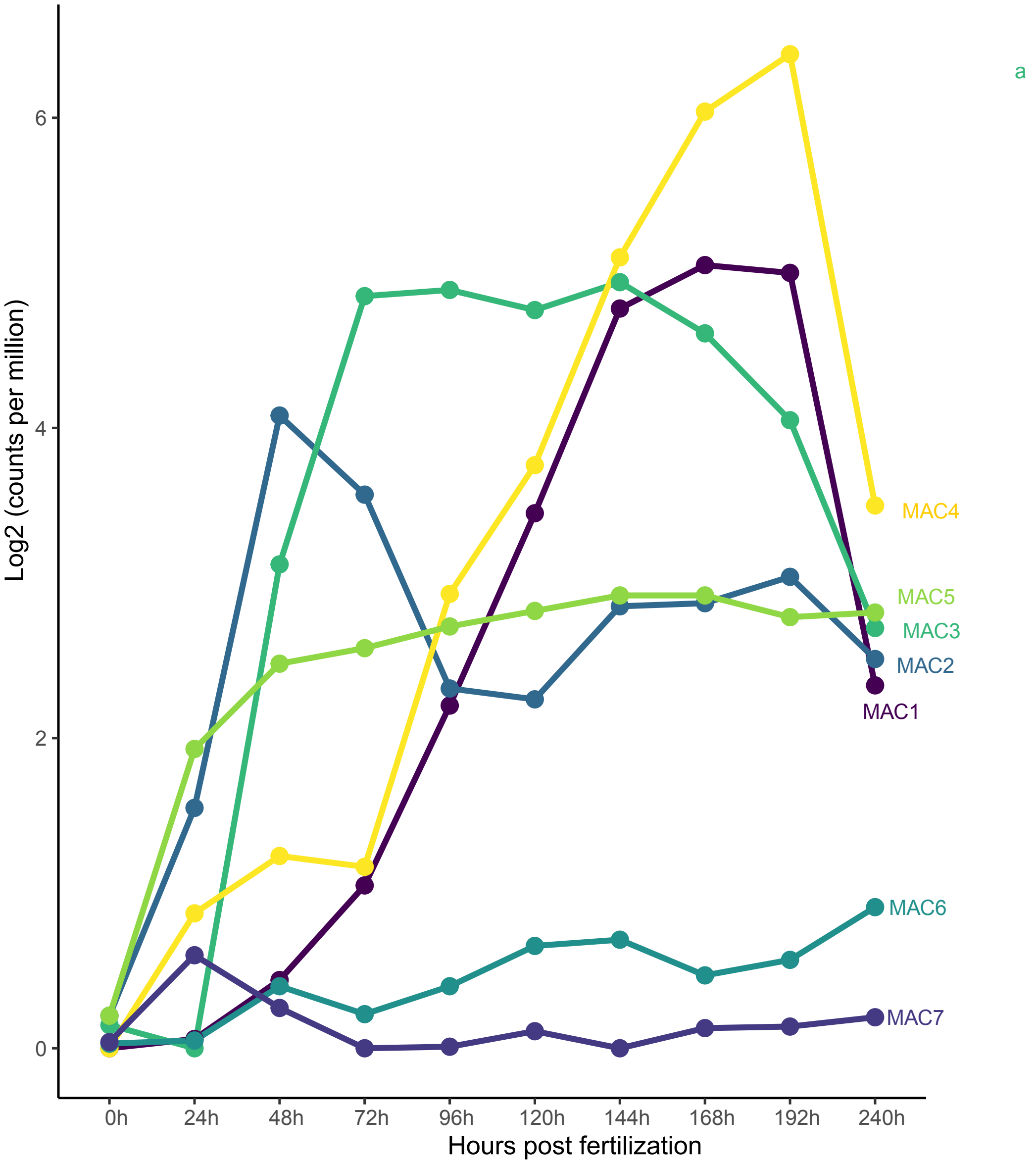


**Fig. S4** Graph of RNA levels of seven MACPF genes throughout embryonic development of Nematostella taken from the NvERTx database

**Table S1.**

Meta analysis and scRNA-seq and bulk RNA-seq across Anthozoa

**Table S2.**

Genomic organization of MACPFs in from Anthozoan genomes and macrosynteny with *Nematostella* and *Scolanthus* chromosomes.

**Table S3.**

Endo-atlas expression of *Nematostella* toxins and MACPFs.

**TableS4.**
nCounter gene expression of NveMACs and housekeeping genes (HKG) across development for both raw (A) and normalized (B).

**TableS5.**
RNA-seq gene expression of NveMACs across development as hours post fertilization (hpf) generated from NvERTX

**TableS6.**
Knockdown efficiency of shRNA and their NveMAC target as well as combined knockdown of NveMAC1, 3 and 4.

**TableS7.**
Settlement ratio of 10dpf *Nematostella* polyps using a shRNA targeting NveMACs 1-4.

**Table S8.**
Downregulation of NveMACs generated using RNA-seq following knockout of HOX genes in *Nematostella* that regulate endomesodermal segmentation.
